# The Smc5/6 complex is a DNA loop extruding motor

**DOI:** 10.1101/2022.05.13.491800

**Authors:** Biswajit Pradhan, Takaharu Kanno, Miki Umeda Igarashi, Martin Dieter Baaske, Jan Siu Kei Wong, Kristian Jeppsson, Camilla Björkegren, Eugene Kim

**Affiliations:** Max Planck Institute of Biophysics, 60438 Frankfurt am Main, Germany; Karolinska Institutet, Department of Cell and Molecular Biology; Biomedicum, Tomtebodavägen 16, 171 77, Stockholm, Sweden; Karolinska Institutet, Department of Biosciences and Nutrition; Neo, Hälsovägen 7c, 141 83, Huddinge, Sweden

**Author notes:** These authors contributed equally to this work (Biswajit Pradhan, Takaharu Kanno).

## Abstract

Structural Maintenance of Chromosomes (SMC) protein complexes are essential for the spatial organization of chromosomes. While cohesin and condensin organize chromosomes by extruding DNA loops, the molecular functions of the third eukaryotic SMC complex, Smc5/6, remain largely unknown. Using single-molecule imaging, we reveal that Smc5/6 forms DNA loops by extrusion. Upon ATP-hydrolysis, Smc5/6 symmetrically reels DNA into loops at a force-dependent rate of 1 kilobase pairs per second. Smc5/6 extrudes loops in the form of a dimer, while monomeric Smc5/6 unidirectionally translocate along DNA. We also find that Nse5 and Nse6 (Nse5/6) subunits act as negative regulators of Smc5/6-mediated loop initiation and stability. Our findings reveal Smc5/6’s molecular functions, and establish loop extrusion as a conserved mechanism among eukaryotic SMC complexes.

**One-Sentence Summary:** Smc5/6 is a DNA-loop-extruding motor, establishing loop extrusion as a conserved mechanism among eukaryotic SMC complexes.

---

The SMC complexes, such as condensin, cohesin and the Smc5/6 complex (Smc5/6), control chromosome organization and regulate most genomic processes, including gene expression, chromosome segregation, and DNA repair (*1*). These multi-subunit complexes are composed of a characteristic ring-shaped trimeric structure containing a pair of SMC ATPases and a kleisin protein, as well as additional regulatory subunits (*2*). Cohesin folds interphase chromosomes into chromatin loops and topologically associated domains (TADs) (*3*–*6*), while condensin organizes mitotic chromosomes in the form of hierarchically nested loops (*7, 8*). Single-molecule experiments have shown that these condensin-and cohesin-mediated DNA loops are formed by an active extrusion process (*9*–*11*). However, whether loop extrusion is a conserved feature of all SMC complexes or specific to condensin and cohesin remains an open question.

Unlike condensin and cohesin, the functions of the third eukaryotic SMC complex, Smc5/6, are much less known. Smc5/6 has been implicated in repair of DNA damage by homologous recombination (*12, 13*), in the promotion of chromosome segregation (*14, 15*) and in replication fork stability and progression (*16, 17*). At the molecular level, different modes of action have been suggested, including DNA-DNA tethering (*16, 18*), DNA compaction through direct interactions between multiple complexes or loop extrusion (*19*), and efficient recognition and stabilization of supercoiled and catenated DNA (*14, 19, 20*). In the light of its structural similarities with condensin and cohesin, it seems reasonable to predict that Smc5/6 also performs DNA loop extrusion and/or translocation. However, Smc5/6 also contains complex-specific features that might prevent such activities, making the prediction more uncertain (*21*–*23*).

Here, we therefore isolated *Saccharomyces cerevisiae* (*S. cerevisiae*) Smc5/6 to examine its DNA loop extrusion activity. Size-exclusion chromatography confirmed that the isolate contained the wild-type octameric complex, with all subunits present at a ∼1:1 stoichiometry (**Fig. 1A, S1A-B)**. The complex displayed DNA-stimulated ATPase activity, with a maximum rate of hydrolysis of 1.9 molecules/s (95% confidence interval, 1.7-2.2 molecules/s) (**Fig. 1B, S1C)**, similar to previously recorded activity ranges of Smc5/6 and other SMC complexes (*10, 11, 19, 20, 24*). As expected, no ATP hydrolysis was detected for complexes in which both Smc5 and Smc6 were mutated to prevent ATP binding (KE mutants) or block ATP hydrolysis (EQ mutants) (**Fig. 1C**). We then tested the activity of Smc5/6 by using a single-molecule assay which allows for direct visualization of loop extrusion mediated by SMC complexes (*9, 10, 25*) (**Fig. 1D, S2A-D**). First, both ends of linear 48.5-kilobase pair (kbp) λ-DNA molecules were tethered to a passivated glass surface and stained with Sytox Orange (SxO). Then, DNA molecules were stretched by buffer flow perpendicular to the DNA axes and imaged by total internal reflection microscopy. Upon addition of Smc5/6 and ATP under constant buffer flow, we observed that DNA was initially concentrated into one spot and then gradually extended into an elongating loop (**Fig. 1E, S3A, and movie S1**). We observed loop formation on the majority of DNA molecules (78 %, N_tot_=233 for 2 nM of Smc5/6 and duration of 1000 s). Looping events were also observed in the absence of buffer flow as a loosely compacted DNA punctum which increases in size over time (**Fig. 1F, movie S2**). Application of buffer flow after the maturation of the DNA punctum further verified it as a single loop (**Fig. S3B**). Fluorescence intensity kymographs of DNA (**Fig. 1G**) and the corresponding estimation of the length of DNA within the loop (I_loop_) and outside of the loop (I_up_, I_down_) (**Fig. 1H, S2E-F, S4A**) showed a progressive growth of the loop (with an average loop size of ∼16 kbp, N_tot_=100, **Fig. S5A**), at the expense of DNA outside of the loop until reaching a plateau. Once extrusion was halted, the loops occasionally moved along DNA in both directions (**Fig. S3C, D**) and were finally released (71%, N_tot_=202), either spontaneously in a single step (e.g. **Fig. 1F, G**, 39 %, N_tot_=202) or by gradual shrinking of loops (**Fig. 1I, J**, 32 %, N_tot_=202). We observed no DNA looping in the absence of ATP, in the presence of AMP-PNP, or when the wild-type complex was replaced by ATP binding (KE)– or ATP hydrolysis (EQ)–deficient mutants (**Fig. 1K**). Together, this demonstrates that Smc5/6 can form loops in an ATP hydrolysis dependent manner by actively extruding DNA.

**Figure 1.**
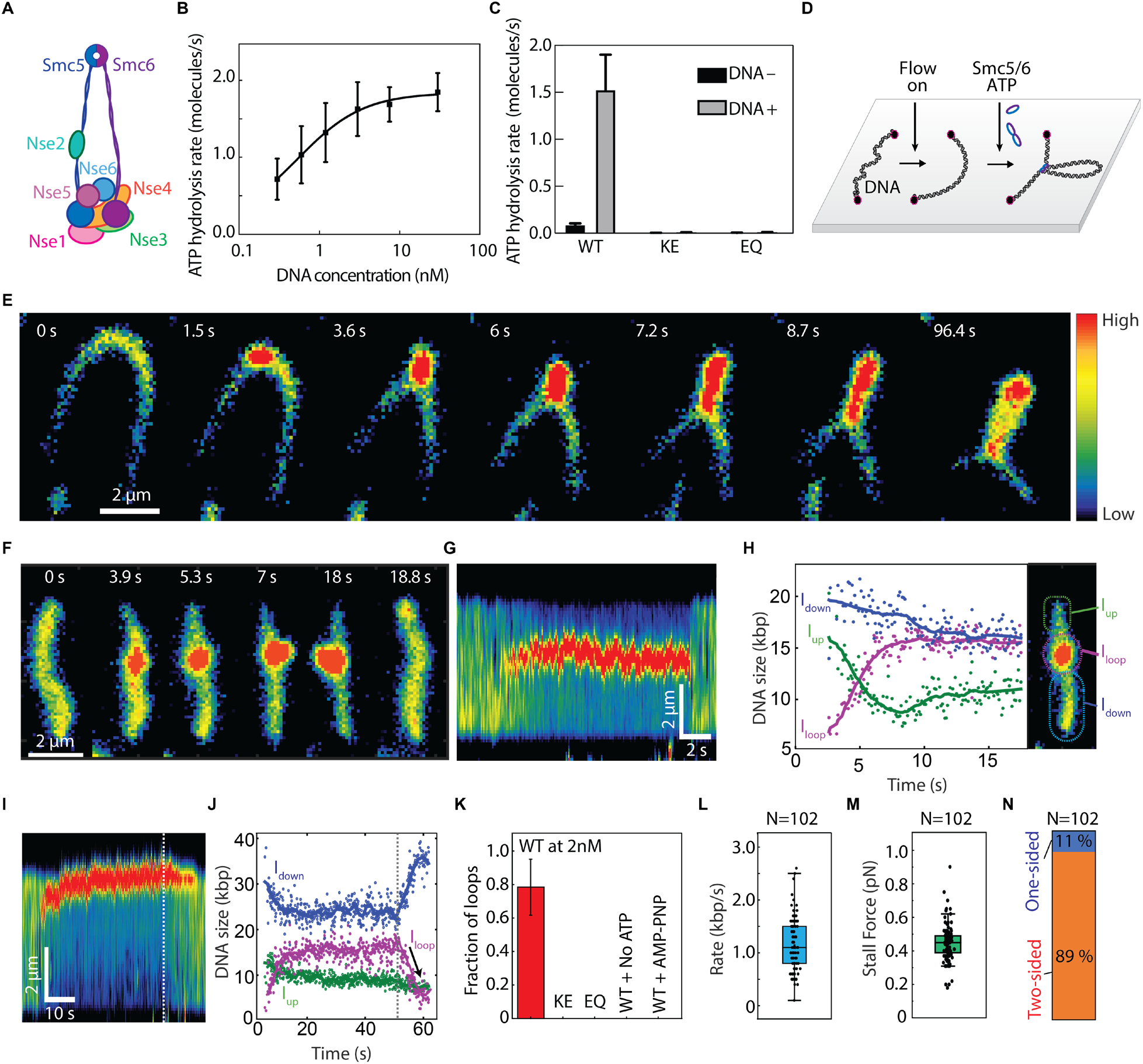
Real-time imaging of loop extrusion by the Smc5/6 complex. (**A**) Cartoon of the *S. cerevisiae* Smc5/6 octameric structure (**B**) ATPase activity of the wild-type Smc5/6 octameric complex in the presence of increasing concentrations of plasmid DNA. Experimental data were fitted to a stimulatory dose-response model by nonlinear regression. N=4, mean ± SD indicated. (**C**) ATPase activity of wild-type, ATP binding–deficient (KE), and ATP hydrolysis–deficient (EQ) Smc5/6 complexes in the absence or presence of 30 nM plasmid DNA. N=3, mean ± SD indicated. (**D**) Schematic of DNA loop extrusion assay. (**E**) Series of snapshots showing DNA loop extrusion intermediates induced by Smc5/6 complex under constant buffer flow. (**F**) Snapshots and (**G**) fluorescence intensity kymograph of a DNA molecule showing DNA loop extrusion in the absence of buffer flow. (**H**) DNA lengths calculated from the kymograph in (G) for the regions outside the loop (I_up_ and I_down_) and the loop region itself (I_loop_). (**I**) Kymograph and (**J**) calculated DNA lengths for a loop extrusion event loop-release by gradual shrinkage. Dashed lines in (I, J) indicate the start of the loop shrinkage. (**K**) DNA loop-forming fractions (mean ± SD from three independent experiments) in the presence of ATP and 2nM Smc5/6 wild-type (WT) and ATPase mutant complexes as in (C), and Smc5/6 in the absence of ATP or presence of AMP-PNP. N=233, 121, 93, 84, and 106, respectively. (**L-M**) Box-whisker-plots of Smc5/6-dependent loop extrusion showing (**L**) rates and (**M**) stalling force. N=102 molecules. (**N**) Fraction of loop extrusion events exhibiting two-sided or one-sided DNA-reeling as determined by observation of DNA length decrease in non-loop regions e.g. I_up_ and I_down_ in (H, J).

We next estimated the speed of loop extrusion (**Fig. 1L**, N_tot_=102) from the initial slopes of the loop growth curves (e.g. **Fig. 1H, S4A-B**), which yielded a rate of 1.1 ± 0.5 kbp/s, a value similar to the reported rates for human cohesin (0.5**–**1 kbp/s (*10, 11*)) and yeast condensin (0.6 kbp/s (*9*)). Loop extrusion by Smc5/6 complex was force-sensitive (**Fig. S4C-F**), again similar to condensin and cohesin. We observed minimal loop formation (∼ 6%) when DNA was stretched above 60 % of its contour length, with a corresponding force of ∼ 0.5 pN (**Fig. S5B, C**). The average stalling force, estimated from the value of relative DNA extension at which loop extrusion was halted and converted to the known force–extension relation (*9*), again yielded 0.5 ± 0.1 pN (**Fig. 1M**). This was close to the values reported for condensin (0.5 pN (*26*)) and cohesin (<0.8 pN (*11*)). The majority of Smc5/6-mediated loop extrusion (89%, N_tot_=102 molecules, **Fig. 1N**) was symmetrical, or ‘two-sided’, as we observed that the DNA length decreased on both sides of the loop in the flow-stretched imaging (e.g. **Fig 1E**), and in the estimated DNA length (I_up_, I_down_) in the absence of flow (e.g. **Fig. 1H**). In summary, the characteristics of loop extrusion mediated by Smc5/6 closely resemble those previously observed for both cohesin and condensin, and is most similar to cohesin which also performs two-sided extrusion.

We then labelled SNAP-tagged Smc5/6 complexes (**Fig. S1D**) with single Alexa647 fluorophores (labeling efficiency = 68 ± 10 %, see Methods) and co-imaged them with DNA during loop extrusion. As expected, this revealed that the complex was positioned at the base of the extruded loop, further confirming an active extrusion process (**Fig. 2A, movie S3**). To determine how many Smc5/6 complexes are required for extrusion, we monitored the fluorescence intensity of labeled loop-extruding complexes in real-time (**Fig. 2B–G**). For the majority of cases (82%, N_tot_= 191), the Smc5/6 signal atop DNA first increased in a single step, indicative of a Smc5/6-DNA binding event, which was followed by loop growth, and finally decreased in either a single or two consecutive steps (**Fig. 2D, G, movie S4, Fig. S6**) due to photobleaching (**Fig. S7**). A smaller fraction (18%) of loop initiation events did not correlate with Smc5/6 signal, indicating looping by unlabeled complexes. The comparisons of intensity distributions obtained from two- and one-step bleached events and background traces confirm that two-step bleaching process originate from two fluorophore-labeled complexes and not more (**Fig. 2H**). Interestingly, we observed a larger fraction (43 %) of two-step bleaching events as compared to single-step bleaching events (36 %) (**Fig. 2I, left hand panel**). Since the labelling-efficiency of Smc5/6 was below 100%, the correlation between the number of bleaching steps (1 or 2) and the number of Smc5/6 molecules (monomer or dimer) is not linear. Importantly, a single bleaching step could arise either from a single labelled Smc5/6 or a Smc5/6 dimer within which only one of the complexes are labelled. We therefore calculated the probability to observe zero (unlabeled), 1, and 2 bleaching steps for the labeling efficiency range of 68 ±10 % as a function of ‘dimer fraction’, where ‘0’ indicates that all Smc5/6 molecules are monomers and ‘1’ indicates all are dimers (**Fig. 2I, right hand panel**, Methods). Interestingly, we find that the observed ratio most closely correlates with a dimer fraction near 100%, indicating that the majority of observed loop extrusion events is performed by Smc5/6 dimers. Real-time imaging of loop extrusion with labeled Smc5/6 under constant buffer flow (**Fig. S8**) revealed that these dimers were located at the stem of the loop during extrusion. We then questioned whether the dimeric state of Smc5/6 is necessary for loop extrusion or if a single complex can extrude a loop, but a second complex is frequently present due to a high likelihood of random complex-complex interaction. If so, the fraction of loop-extruding dimers is expected to decrease with decreasing protein concentrations. Interestingly, however, we observed that the fraction of two bleaching steps did not decrease even at 10 times lower protein concentration but instead remained consistently larger than the fraction of single bleaching steps (**Fig. 2J**). Furthermore, fitting the fraction of looped DNA observed at different Smc5/6 concentrations to a Hill–Langmuir equation showed that loop extrusion is stimulated by cooperative interactions, displaying a Hill-factor of *n* = 1.84, i.e. well in excess of 1, which indicates the absence of cooperativity (**Fig. 2K**, Methods). All together, these observations support that the functional unit for Smc5/6-dependent loop extrusion is a dimer of complexes. This contrasts with condensin, which extrudes loops as a single complex (*24*), whereas cohesin has been suggested to extrude both as a monomer and dimer (*10, 11*).

**Figure 2.**
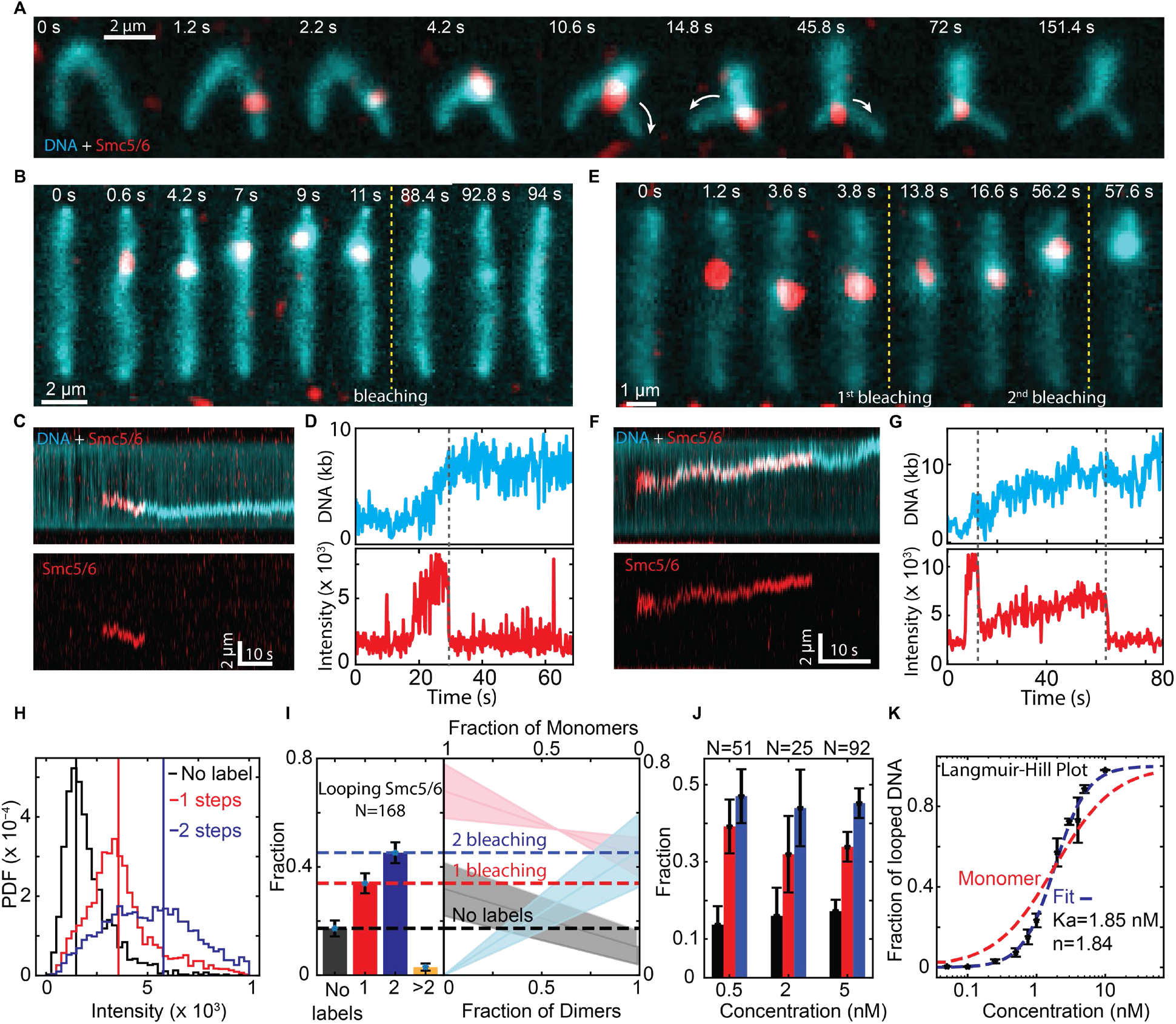
Dimers of Smc5/6 complexes extrude DNA loops. Snapshots of image overlays showing SxO stained DNA (cyan) and Alexa647 labeled Smc5/6 complexes (red) during loop extrusion in the presence (**A**) and absence (**B, E**) of constant buffer flow, and exhibiting one (**B**) and two (**E**) photobleaching events. Arrows in (A) indicate the direction of the Smc5/6 movement. (**C, F**) Kymographs of the two loop extrusion events in (**B, E**) depicting the intensity overlays of DNA and Smc5/6 (top) and only Smc5/6 (bottom). (**D, G**) Time traces of DNA length (top) and Smc5/6 fluorescence intensity (bottom) determined from (**C, F**) with bleaching events indicated as dashed vertical lines. (**H**) Probability density functions (PDF) of fluorescence intensities for loop extrusion events exhibiting no Alexa647 signal, or one- or two-step bleaching. (**I**, left panel) Fraction of loop extruding Smc5/6 events that displayed either no, one, two or more bleaching steps. If loop formation was interrupted before bleaching occurred, the number of participating Smc5/6 complexes was estimated from the intensity. (**I**, right panel) The probabilities for finding either 0 (gray shaded area), 1 (red shaded area) or 2 (blue shaded area) labels as a function of Smc/6 dimer over monomer ratio estimated on basis of the labeling efficiency of 68 ± 10%. (**J**) Histogram showing the number of bleaching steps (0: black, 1: red, 2: blue) observed during individual loop extrusion events at indicated Smc5/6 concentrations. (**K**) Hill–Langmuir plot showing the fraction of DNA substrates that formed loops as a function of the Smc5/6 concentration (black dots). The respective fit (blue dashed line) indicates cooperative behavior (n=1.84), deviating from the Hill-Langmuir function expected for exclusively monomeric loop extrusion (n=1, red dashed line). Experiments were performed using the wild-type octameric complex and with 10s durations.

In addition to loop extrusion, we also observe that Smc5/6 can unidirectionally translocate along DNA in an ATP-dependent manner (**Fig. 3A, movie S5, Fig. S9A-B, S10C**), again mirroring condensin (*27*) and cohesin (*10*). Kymographs revealed that ∼80% (N_tot_=73) of all non-loop-extruding Smc5/6 translocated, while a smaller fraction of molecules remained either stably bound at one position, or randomly diffused along DNA (**Fig. 3B, C, S10A-B, S10D-E**). Mean squared displacement (MSD) plots generated from tracking labelled Smc5/6 in kymographs exhibit increasing slopes, consistent with directed motion with an average translocation velocity of 1.5 ±0.2 kbp/s (**Fig. 3B**, N_tot_=32). The photobleaching steps and the fluorescence intensity distribution of translocating Smc5/6 (**Fig. 3D, E**) revealed that ∼92% of translocating units were labeled with a single fluorophore. By comparing this value with calculated observation probability of single bleaching steps (**Fig. S9C**), we find that our data matches closest with a fraction of monomers larger than 90%. Together, these findings indicate that a single Smc5/6 can translocate along DNA whereas a pair of complexes is required for DNA loop extrusion. In further support of this, 4% of all loop extrusion events were initiated when second Smc5/6 complex associated to a single translocating complex **(Fig. 3F, G, movie S6, Fig. S11**).

**Figure 3.**
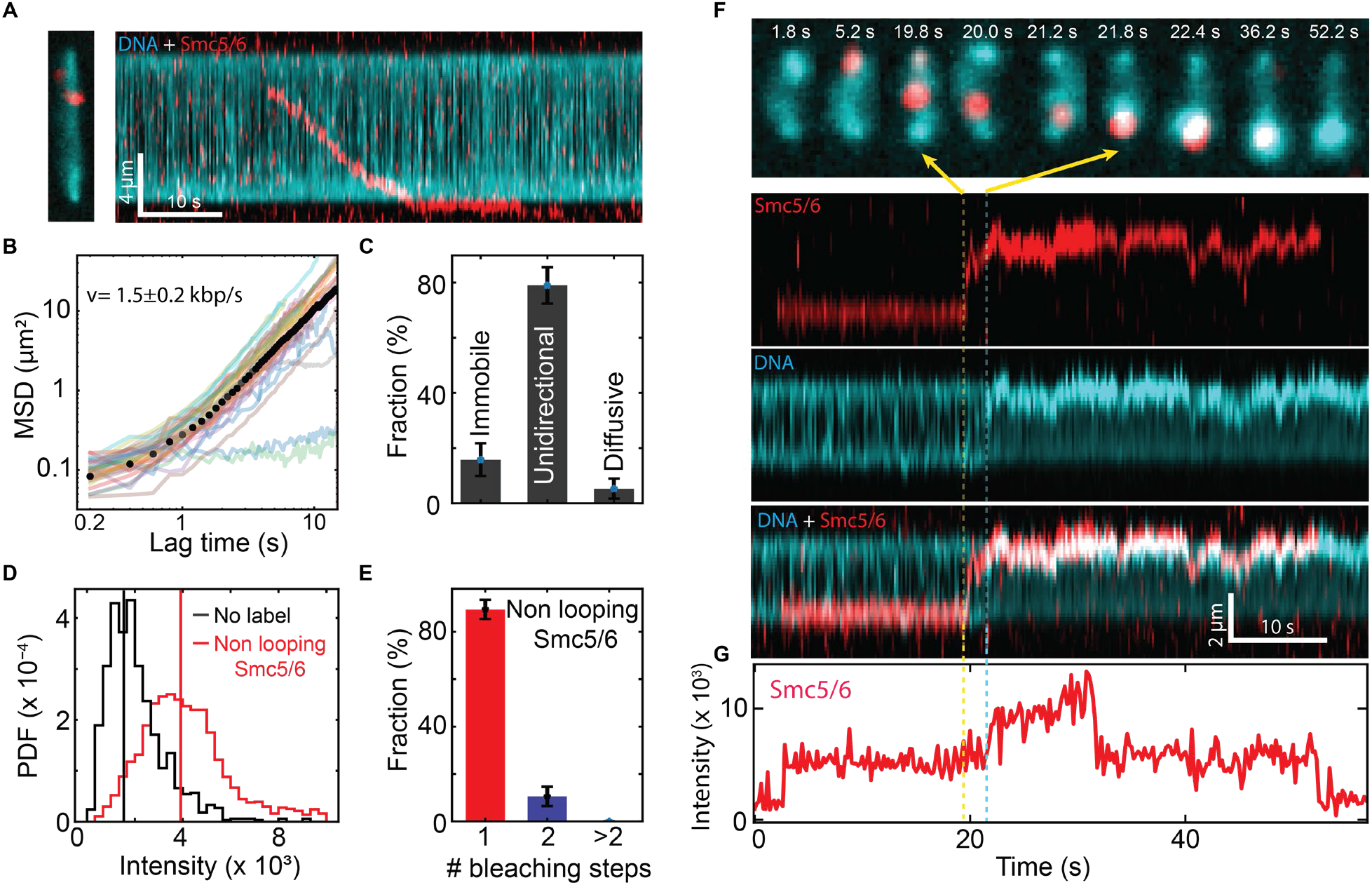
Single Smc5/6 complexes unidirectionally translocate along DNA. **(A)** Overlaid snapshot (left) and kymograph (right) showing an example for a labeled Smc5/6 complex translocating on a DNA molecule. (**B**) Mean squared displacement (MSD) plots determined from kymograph-trajectories of Smc5/6 translocation events (solid lines; Average: dotted line). (**C**) Fractions of non-loop extruding Smc5/6 complexes that remained immobile or translocated directional or diffusive on DNA before their dissociation from the DNA. (**D**) Probability density functions (PDF) for fluorescence intensities of background and non-looping Smc5/6 complexes. (**E**) Fractions of DNA-bound and non-looping Smc5/6 complexes that exhibited 1 (red), 2 (blue) or more bleaching steps. (**F**) Kymographs of labeled Smc5/6 (top), DNA (center) and its overlay (bottom) and the corresponding time trace (**G**) of labeled Smc5/6 intensity showing an event where a translocating Smc5/6 starts extruding a DNA loop upon forming a dimer with another Smc5/6 from solution.

Next, we investigated the regulatory role of the Nse5/6 subunits in the loop extrusion process. Previous studies (*21, 28*) showed that Nse5/6 negatively regulates the ATPase activity of Smc5/6 in the absence of DNA, while in the presence of DNA this negative effect is abolished. To test whether and how this is reflected in the process of loop extrusion, we purified the hexametric complex which lacks Nse5/6, and confirmed the elevated basal ATPase activity without DNA (**Fig. 4A, S12**). Subsequent loop extrusion experiments revealed that the hexamer extrudes loops at a rate of 1.3 ± 0.6 kbp/s, similar to the rate of the wild-type octameric complex (**Fig. 4B**). In contrast to the unchanged extrusion rate, the probability to observe loops was ∼ 12 times higher for the hexamer as compared to the wild-type octamer (**Fig. 4C**) at 0.5 nM protein concentration. The stability of loops also increased in the absence of Nse5/6 as we observed ∼2.7 times less loop release events within 1000 seconds after initial loop formation. (**Fig. 4D**). In summary, our data shows that Nse5/6 regulates loop extrusion by reducing loop initiation rate and loop persistence. When eventually initiated, Nse5/6 have little effect on the extrusion process as evidenced by the unchanged extrusion speed. Together with our finding that Smc5/6 preferentially extrudes loops as a dimer, we speculate that the reduction in loop initiation and stability mediated by Nse5/6 might be due to negative regulation of complex dimerization. Intriguingly, cryo-EM analysis of the hexamer have indeed revealed dimers of complexes interacting via the ATPase-containing so-called head domains (*28*).

**Figure 4.**
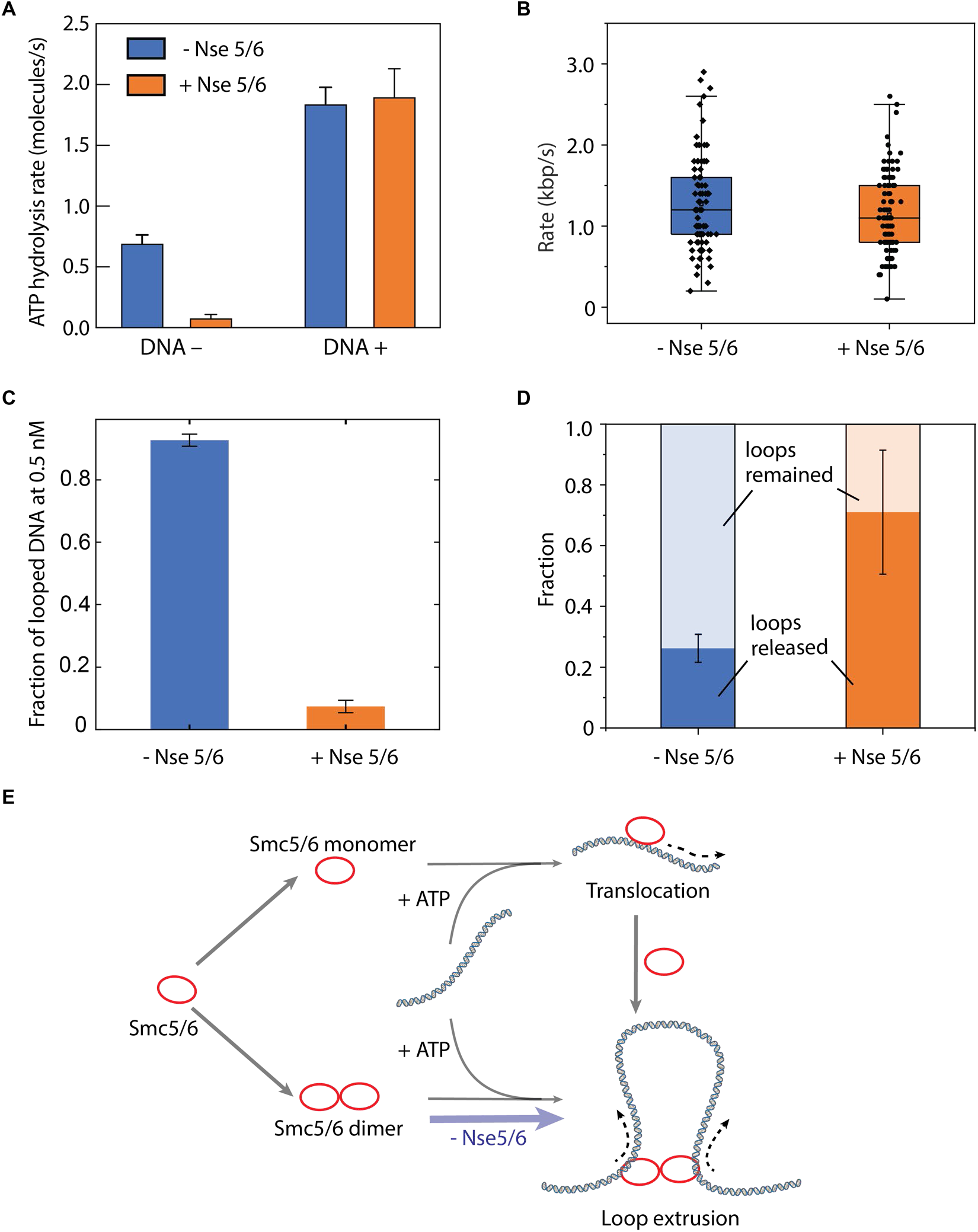
The Nse5/6 subcomplex is a negative regulator of DNA loop extrusion. (**A**) ATPase rate of the hexamer (-Nse5/6) and the wild-type, octameric complex (+Nse5/6) in the absence and presence of 30 nM plasmid DNA. Data for the wild-type complex is identical to Fig. 1B and shown for comparison. N=3, mean ± SD indicated. (**B**) Box-whisker-plots of loop extrusion rates of the hexameric (-Nse5/6) and octameric (+Nse5/6) complexes. N= 82 and 102 molecules, respectively. (**C**) Fraction of DNA molecules (mean ± SD from three independent experiments) that formed loops upon addition of 0.5 nM hexameric (-Nse5/6) and wild-type complexes(+Nse5/6). N= 228 and 236 molecules. (**D**) Fraction of loops (mean ± SD from three independent experiments) which were either released (dark blue/orange) or maintained (light blue/orange) during a measurement period of 1000s for hexameric (-Nse5/6) and wild-type octameric (+Nse5/6) complexes, respectively. N= 245 and 202 events. (**E**) Model of Smc5/6 loop extrusion. Dimeric Smc5/6 performs two-sided loop-extrusion, while a monomeric Smc5/6 translocates unidirectionally along DNA. Nse5/6 negatively regulates loop initiation and maintenance.

In conclusion, our study shows that Smc5/6 performs loop extrusion in a manner that is functionally similar to cohesin and condensin. This said, Smc5/6-mediated loop extrusion also exhibits specific characteristics, namely, is preferentially executed by cooperative pairs of complexes, while single complexes translocate along DNA (**Fig. 4E**). We also find that loop extrusion is down-regulated by the Nse5/6 subunits. Taken together, our findings indicate that Smc5/6 protects genome integrity using active extrusion, and open up new avenues for dissecting the activities of this multifunctional complex.

## Acknowledgments

we are grateful to J. Diffley and colleagues for sharing plasmids for inducible over-expression. We thank J. van der Torre for the protocol for biotinylating λ-DNA.

## Author contributions

Project initiation: CB, EK Funding acquisition: CB, EK

Development of investigation: BP, TK, KJ, CB, EK Smc5/6 purification and labelling: TK, MUI Smc5/6 ATPase analysis: MUI

Single molecule experiments: BP, EK

Single Molecule analysis: BP, EK, JSKW, MDB Building single molecule instrumentation: BP, MDB Supervision: CB, TK, KJ, EK

Writing original draft: CB, EK

Writing – review & editing: BP, TK, MUI, KJ, MDB, JSKW, CB, EK

## Data and materials availability

Original imaging data and protein expression constructs are available upon request.

Supplementary Text Figs. S1 to S12 Tables S1 to S2

References

Captions for Movie S1 to S6

