## Supplementary Materials for "The Smc5/6 complex is a DNA loop extruding motor"

**This PDF file includes:**

Table S1, S2

Materials and Methods

Figs. S1 to S12

Captions for Movie S1 to S6

**Other Supplementary Materials for this manuscript include the following:**

Movie S1, S2, S3, S4, S5, S6

**Table S1. Yeast strains used in this study.**

All strains are of W303 origin, *MATa, ade2-1, trp1-1, can1-100, leu2-3, leu112, his3-11, 15, ura3 and RAD5*, with the modifications listed below.

| *Strain* | *Modification* |
|---|---|
| CB3245 | *MATa pep4Δ::HPH* |
| CB3573 | *MATa pep4Δ::HPH top1Δ::NAT*<br>*leu2::LEU2pRS305-NSE1-pGAL1-10-NSE2*<br>*his3::HIS3pRS303-NSE3-pGAL1-10-NSE4*<br>*ura3::URA3pRS306-SMC5-pGAL1-10-SMC6-TAP* |
| CB3580 | *MATa pep4Δ::HPH top1Δ::NAT*<br>*trp1::TRP1pRS304-NSE5-pGAL1-10-NSE6*<br>*leu2::LEU2pRS305-NSE1-pGAL1-10-NSE2*<br>*his3::HIS3pRS303-NSE3-pGAL1-10-NSE4*<br>*ura3::URA3pRS306-SMC5-pGAL1-10-SMC6-TAP* |
| CB3662 | *MATa pep4Δ::HPH top1Δ::NAT*<br>*trp1::TRP1pRS304-NSE5-pGAL1-10-NSE6*<br>*leu2::LEU2pRS305-NSE1-pGAL1-10-NSE2*<br>*his3::HIS3pRS303-NSE3-pGAL1-10-NSE4*<br>*ura3::URA3pRS306-smc5K75E-pGAL1-10-smc6K115E-TAP* |
| CB3663 | *MATa pep4Δ::HPH top1Δ::NAT*<br>*trp1::TRP1pRS304-NSE5-pGAL1-10-NSE6*<br>*leu2::LEU2pRS305-NSE1-pGAL1-10-NSE2* |

| | |
|---|---|
| | *his3::HIS3pRS303-NSE3-pGAL1-10-NSE4*<br><br>*ura3::URA3pRS306-smc5E1015Q-pGAL1-10-smc6E1048Q-TAP* |
| CB4005 | *MATa pep4Δ::HPH top1Δ::NAT*<br><br>*trp1::TRP1pRS304-NSE5-pGAL1-10-NSE6*<br><br>*leu2::LEU2pRS305-NSE1-pGAL1-10-NSE2*<br><br>*his3::HIS3pRS303-NSE3-pGAL1-10-NSE4-6xHis-SNAP-KAN*<br><br>*ura3::URA3pRS306* |

**Table S2. Plasmid DNA used in this study.**

| *Name* | *Description* | *Source* |
|---|---|---|
| pJF2 | pRS303::*CDT1-pGAL1-10-GAL4* | Gift from Prof. John Diffley[1] |
| pJF3 | pRS304::*MCM4-pGAL1-10-MCM5* | Gift from Prof. John Diffley[1] |
| pJF4 | pRS305::*MCM6-pGAL1-10-MCM7* | Gift from Prof. John Diffley[1] |
| pJF5 | pRS306::*MCM2-pGAL1-10-MCM3* | Gift from Prof. John Diffley[1] |
| CD373 | pRS306-*SMC5-pGAL1-10-SMC6-TAP* | This study |
| CD377 | pRS30-*NSE1-pGAL1-10-NSE2* | This study |
| CD380 | pRS304-*NSE5-pGAL1-10-NSE6* | This study |
| CD395 | pRS303-*NSE3-pGAL1-10-NSE4* | This study |
| CD402 | pFA6a-SNAP-KAN | This study |
| CD406 | pRS306-*smc5K75E-pGAL1-10-smc6K115E-TAP* | This study |
| CD420 | pRS306-*smc5E1015Q-pGAL1-10-smc6E1048Q-TAP* | This study |

***Materials and Methods***

**Gene synthesis, sub-cloning and strain creation for Smc5/6 overexpression.** Genes for all Smc5/6 subunits were synthesized by GeneArt Gene Synthesis (Thermo Fisher Scientific) with codon optimization and introduced into pJF2, pJF3, pJF4 and pJF5 yeast integrative vectors (kindly provided by Prof. John Diffley) under bidirectional *GAL1-10* promoter *(29)*. The TAP-tag sequence derived from pBS1479 (EUROSCARF) was introduced into the C-terminus of Smc6 using standard methods. The following plasmids, CD373; *SMC5-GAL1-10-SMC6-TAP*, CD380; *NSE5-pGAL1-10-NSE6*, CD395; *NSE3-pGAL1-10-NSE4*, CD377; *NSE1-pGAL1-10-NSE2*, were integrated into CB3245 using auxotrophic markers, then the *TOP1* gene was deleted using standard gene-replacement methods. For ATPase mutants, point mutations were introduced using standard methods at appropriate positions (Walker A (KE); Smc5K75E, Smc6K115E, Walker B (EQ); Smc5E1015Q, Smc6E1048Q). Codon-optimized SNAP tag was synthesized by GeneArt Gene Synthesis and introduced in pFA6a with KAN marker. The SNAP tag was introduced into the C-terminus of ectopic NSE4 using standard methods.

**Overexpression and purification of Smc5/6.** Overexpression strains were grown at 30°C in 1 or 2 l of YEP-lactate medium to $OD_{600:}$: 0.8-1.0, then protein expression was induced for 4 hours by addition of 2% galactose addition of one cell volume of IPP150 buffer (50 mM Tris-HCl [pH 8.0], 150 mM NaCl, 10% glycerol, 0.1% IGEPAL CA-630, 1 mM dithiothreitol (DTT)) containing 10 mM $MgCl_2$ and complete EDTA free protease inhibitor (Roche Applied Science), whereafter treatment with benzonase (Merck) was performed for 1 h at 4ºC. Cleared extracts were mixed with IgG Sepharose 6 FF (Merck) for 2 h at 4ºC and washed with IPP150 buffer. Then IPP150 buffer was replaced with GF500 buffer (20 mM HEPES-NaOH [pH 7.5], 500 mM NaCl, 10% Glycerol, 0.1% IGEPAL CA-630, 1 mM DTT), and the resin was treated with tobacco etch virus (TEV) protease (kind gift from Dr. Herwig Schüler) at 4ºC overnight. The fraction eluted by TEV treatment was diluted four-fold in CBB500 buffer (50 mM Tris-HCl [pH 8.0], 500 mM NaCl, 1 mM $Mg(CH_3COO)_2$, 1 mM imidazole, 2 mM $CaCl_2$, 1 mM DTT, 0.1% IGEPAL CA-630), supplemented with 0.75 mM $CaCl_2$, and incubated with calmodulin Sepharose 4B (Merck) for 2 h at 4ºC. After washing with CBB500 buffer, proteins were eluted using CEB500 buffer (50 mM. Purification of Smc5/6 was performed according to the method published previously with the following modifications *(29).* Briefly, cells were disrupted using Freezer mill 6870 (SPEX), and proteins extracted by Tris-HCl [pH 8.0], 500 mM NaCl, 1 mM $Mg(CH_3COO)_2$, 1 mM imidazole, 20 mM EGTA, 1 mM DTT, 0.1% IGEPAL CA-630). The eluate was concentrated ~50-fold using Vivaspin20 ultrafiltration unit (100K MWCO; Sartorius) concomitant with an exchange to STO500 buffer (50 mM Tris-HCl [pH 8.0], 500 mM NaCl, 2 mM $MgCl_2$, 0.5 mM tris(2-carboxyethyl)phosphine (TCEP), 10% Glycerol, 0.1 % IGEPAL CA-630). Concentration of the complex was determined by Bradford assay using BSA as standard. The integrity of purified Smc5/6 was tested using size exclusion chromatography on Superose 6 Increase 10/300 GL column (GE healthcare), preequilibrated with STO500 buffer, and subsequent SDS-PAGE analysis of eluted fractions (Fig. S1B for the wild-type octameric complex, and S12B for a hexamer lacking the Nse5 and Nse6 subunits).

**Fluorescent labelling of Smc5/6.** Using the CB4005 strain (Table S1) an Smc5/6 complex containing C-terminally tagged Nse4-6xHis-SNAP was overexpressed and purified using IgG Sepharose 6 FF as described above. After TEV protease cleavage, the eluate was concentrated ~50-fold using Vivaspin20 ultrafiltration unit (100K MWCO) concomitant with an exchange to STO500 buffer. For fluorescent labelling, the eluate was mixed with 20 μM SNAP-Surface Alexa Flour647 (NEB) in 50 μl of STO500 buffer supplemented with 50 mM DTT and incubated for overnight at 4ºC. The mixture was concentrated ~10-fold using Amicon Ultra centrifugal filter (100K MWCO; Merck) concomitant with the buffer exchange to fresh STO500 buffer for removal of free Alexa Flor647.

**Labelling efficiency estimation**

The labelling efficiency was calculated in two steps. First the amount of Smc5/6 was estimated from Bradford assay using BSA as standard which came to be 7.56 ± 0.5 μM. Then the amount of label (Alexa647) was estimated in by comparing both the absorption and the fluorescence intensity of a known concentration (e.g. 1 μM) of Alexa647 in the same storage buffer as used for labelled Smc5/6. Both absorption and fluorescence measurements yielded same labelling efficiency of 68 ± 10 % within their errors.

**ATPase assay.** Smc5/6 (0.5 μl, final concentration 30 nM) was incubated with 4 nCi [α-$^{32}$P]ATP in 5 μl of the reaction buffer (50 mM Tris-HCl [pH 7.6], 40 mM KCl, 1 mM $MgCl_2$, 1 mM DTT, 0.1 mg/ml BSA) containing 1 mM ATP and various concentration of pRS316 at 30°C. Aliquots (1 μl) were collected every 30 min for 90 min, and mixed with 1.5 μl of 1% SDS to stop the reaction. Then 1 μl of the mixture was spotted on TLC PEI cellulose F plates (MERCK) and developed in 1 M HCOOH/0.5 M LiCl. Radiolabeled ATP and ADP were quantified using a LAS-3000 imager (Fujifilm). ATPase rates at each DNA concentration were calculated by linear regression using least squares method. Maximum ATPase rate and the 95% confidence interval for the wild-type complex was obtained by fitting of a stimulatory dose-response model to experimental data by nonlinear regression using Prism 9 software (GraphPad).

**HiLo microscopy and Data collection**

A custom-built microscope was used for single-molecule visualization of DNA and labelled Smc5/6. Lasers with wavelengths of 638 nm (Cobolt) and 561 nm (Coherent) were coupled to a Zeiss (AxioVert200) microscope body through a single mode fibre in wide-field illumination mode with the possibility of changing the illumination angle. The setup allows us to use highly inclined optical light sheet (HiLo) illumination using a TIRF objective (alpha-Plan-APOCHROMAT 100x/1,46 Oil) in order to selectively image DNA and Smc5/6 while minimizing out of focus fluorescence background and bleaching. The fluorescence signal from the sample was spectrally selected by a dichroic filter (t405/488/561/640rpc2, Chroma) and recorded with a sCMOS (PCO edge 4.2) camera. Light from the excitation lasers (638 nm and 561 nm) was additionally suppressed using a multiband notch filter (NF03-405/488/561/635E-25, Semrock) located before the camera. For simultaneous imaging of DNA and SMC5/6, alternative excitation (ALEX) between the 561 nm and 638 nm laser was used through electronic triggering of an AOTF filter (MPDSnCxx-ed1-18 and AOTFnC_MDS driver from AA-Optoelectronic). The temperature of the flow cell was controlled by adjusting the electric current send through a self-adhesive heating foil (Thermo TECH Polyester Heating foil self-adhesive 12 V DC,

12 V AC 17 W IP rating IPX4 (L x W) 65 mm x 10 mm) attached a top of the glass slide. The temperature was set to $30^0$C for all the experiments unless otherwise stated.

A custom-written python software was used for recording, storing and visualization of data. Specifically, we utilized PyQtGraph (https://github.com/pyqtgraph/pyqtgraph) and napari (https://github.com/napari/napari) for visualization and export of images. Typically, images were recorded with 100-ms exposure time per frame and for duration of 1000-2000 s unless otherwise stated.

**Single-molecule loop extrusion assay**

*Flow cell preparation*

The single-molecule assay used throughout this work was prepared as described before *(9, 25, 30)* with the following slight modifications: Microscope slides were cleaned with acid piranha (sulfuric acid (5parts) and hydrogen peroxide (1 part)) and silanized with 3-(2-aminoethyl)aminopropyl] trimethoxysilane in methanol containing 5% glacial acetic acid which leaves free amine groups on the surface. The slides were then treated with 5 mg/ml methoxy-PEG-N-hydroxysuccinimide (MW 3500, Laysan Bio) and 0.05 mg/ml biotin-PEG-N-hydroxysuccinimide (MW3400, Laysan Bio) in 50 mM Borate buffer, pH 9. The pegylation step was repeated five times to minimize nonspecific surface-sticking of proteins. The pegylated slides were dried with a gentle flow of nitrogen, sealed, and stored at -20$^{o}$ C until further use. Flow cells were then assembled with the functionalized glass slides as described before *(9)*. Each flow cell contains one inlet and two outlet channels in order to allow for buffer flow application perpendicular to the axis of the immobilized DNA. The fluidic channels were first incubated with100 nM streptavidin in

T50 buffer (40 mM Tris-HCl pH 7.5, 50 mM NaCl) for 1 min and then washed thoroughly with T50 buffer. Subsequently, 10 pM of Phage λ DNA molecules, that were labelled with biotins at both of their ends *(31)*, were introduced to the flow cell at a constant speed of 3 μl/min, resulting in the surface-immobilization of DNA molecules with relative DNA extensions ranging from 0.1 to 0.6. Unbound DNA molecules were washed out afterwards. In order to minimize unwanted surface-sticking of Smc5/6, the flow cell was further passivated by incubation with 0.5 mg/mL BSA for 5 min.

*Single molecule imaging of Smc5/6 mediated loop extrusion*

Real-time imaging of Smc5/6 mediated loop extrusion was carried out as following. The imaging buffer (40 mM Tris-HCl pH 7.5, 100 mM NaCl, 7.5 mM MgCl2, 0.5 mg ml-1BSA, 1 mM TCEP, and 2 mM ATP, 200 nM Sytox Orange (SxO), 30 mM D-glucose, 2 mM Trolox, 10 nM Catalase, 37.5 uM Glucose oxidase) containing Smc5/6 (2nM, unless otherwise mentioned) was introduced into the flow cell at flow rate of 30 μL/min for 1 minute and the flow was stopped thereafter. For the side-flow experiment, a larger volume (200-300 ul) of the sample with the same composition was continuously flown into the channel at 15 μL/min. If only SxO-stained DNA was imaged only the 561-nm laser was used at an intensity of 0.1 W/cm$^2$; whereas for dual-color imaging, SxO-stained DNA and Alexa647-labelled Smc5/6 were imaged by alternating excitation using a 531 nm (0.1 W/cm$^2$) and a 638 nm (~150 W/cm$^2$) laser.

**Data analysis**

Fluorescence images were analysed using a custom-written python software (*30–32*). Regions containing λ-DNA molecules were chosen manually and cropped and saved into

TIFF format. For the snapshots of the molecules shown in this paper (e.g. Fig. 1E etc.), the background was subtracted using "white_tophat" filter in scipy *(33)* . For the further quantification of fluorescence intensity, an additional median filter (radius: 2 pixels) was applied. The snapshots of labelled Smc5/6 (e.g. Fig. 2A, B, E) were denoised using a machine-learning based method ("Noise2void") for better visualization. For the further quantification of fluorescence intensity (i.e. for building kymographs), however, the same median filter as used for the DNA was employed (*34*).

From the median filtered images, kymographs were subsequently built by summation of fluorescence intensities over 11 pixels along lines centered around the DNA axis (e.g. Fig. 1G, I). The intensities in the kymograph were normalized such that the values outside of DNA approached zero. Each vertical pixelated line in the kymograph corresponds to one time point (one image-frame) of the image sequence.

*Estimation of DNA size*

The position of the DNA punctum, i.e., the centre position of a DNA loop, in the DNA-kymograph was found by using the "find_peaks" algorithm of scipy, which determines the positions of local peak maxima for each line of the kymograph. Then we select the most intense peak along each line on the kymograph. The intensity of DNA puncta, i.e., the entire region containing the DNA loop was obtained by summing over the area of a square with 7 pixels side length and centred around the punctum position and termed as $Int_{loop}$ (Fig. S2E). The rest of the DNA regions outside of the puncta were separated into two categories, termed as $Int_{up}$ and $Int_{down}$, where 'up/down' is the intensity from the DNA region above/below the puncta (See e.g. Fig. 1H). The amount of DNA in the loop, above and below the loop (Fig. S2E) were estimated by multiplying the fraction of the intensities in the respective regions by 48.5 kb (the length of the used lambda DNA in bp), i.e.

DNA size in the loop (bp), $I_{loop} = \frac{Int_{loop} \times 48502}{Total\ DNA\ intensity}$,

DNA size above the loop (bp), $I_{up} = \frac{Int_{up} \times 48502}{Total\ DNA\ intensity}$

DNA size below the loop (bp), $I_{down} = \frac{Int_{down} \times 48502}{Total\ DNA\ intensity}$

The change of DNA sizes in the respective regions were plotted as a function of time as shown in e.g. Fig. 1H, 1J, Fig. S2F. These data were plotted together with the data smoothed using a Savitzky-Golay filter of 2$^{nd}$ order and window size of 50 points (solid curve, Fig. 1H, 1J). For the Figure 1N, the loop extrusion was termed two-sided if the increase in $I_{loop}$ was correlated with a decrease in both $I_{up}$ and $I_{down}$, otherwise the events were termed as one-sided. To estimate the rate (k) of loop extrusion, the initial 5 seconds of the loop growth curve was fitted (Fig. 1L, Fig. S4B) with a line $I_{loop} = kt + c$ , where c compensates for the initial DNA amount before loop extrusion. Time traces of loop extrusion rates were then obtained using $\mathrm{k(t)} = \left[I_{loop}(\mathrm{t} + \mathrm{dt}) - I_{loop}(t - dt)\right]/2dt$ , with dt = 1 seconds and consequent smoothing with the Savitzky-Golay filter as described above (Fig. S4C). For the force estimation, the relative extension was first calculated as $R_{ext} = 48502\ d/\left(\left(I_{up} + I_{down}\right)L_C\right)$, where $L_C = 16$ µm is the contour length of λ-DNA and d is the end-to-end distance of the double tethered DNA in µm (Fig. S4D-E). Subsequently, the relative extension was converted to force (Fig. S4F) via linear interpolation of the force-extension curve obtained by magnetic tweezer force spectroscopy (*35*). The force at the maximum loop size was taken as the stalling force (Fig. 1M).

*Estimation of number of Smc5/6*

For the determination of the number of Smc5/6 required for loop extrusion (e.g. Fig. 2C, D, F, G), image sequences recorded using ALEX were used to build kymographs of DNA and labeled Smc5/6 intensities. The same areas, which were used to determine intensities of DNA puncta, were used to determine Smc5/6 intensities and build intensity time traces which were then used to count bleaching steps (e.g. Fig. 2D) and to build intensity histograms (e.g. Fig. 2H). For the determination of the photobleaching statistics shown in Fig.2I we only included the molecules for which we observed loop initiation during the recording interval, i.e., loops that were already initiated before the recording started were excluded from analysis. The bleaching times for one-step bleaching ($\Delta\tau_{11}$) and the bleaching times for two-step bleaching ($\Delta\tau_{21}$, $\Delta\tau_{22}$) were than used to calculate the respective average bleaching times and to build the histogram shown in Fig. S7.

In order to quantify the fluorescence intensity of labeled Smc5/6, Smc5/6 complexes which did not perform loop extrusion, were localized and separately categorized as "Non-looping Smc5/6". The intensity trace of Smc5/6 was calculated by summing the intensities over the square centered around the localized position of Smc5/6 in the kymograph. Smc5/6 molecules that bound and were stuck at the end of the DNA near the PEG surface were not considered for further analysis.

The mean square displacement (MSD) of non-looping Smc5/6 molecules were calculated from the traces of their respective positions determined using trackpy (*36*). Positions were tracked until individual Smc5/6 had reached either end of a DNA construct. The MSD (Fig. 3B) was fitted with a directed motion equation: $MSD(t) = v^2t^2 + 4Dt$ where $v =$ mean velocity, $D$ = Diffusion coefficient, and $t$ is the lag time. The velocity in µm/s obtained from these fits was then converted to kbp/s as $v\left(\frac{kbp}{s}\right) = v\left(\frac{\mu m}{s}\right) \times 48.5kbp/L_{avg}$, where $L_{avg} = 9\mu m$ is the average end-to-end distance of the DNAs on which translocation was observed.

**Hill-Langmuir Plot**

Loop extrusion experiments were performed at different concentration of wild-type Smc5/6. These measurements with 10 s duration each were recorded after an incubation period of 15 minutes. At this time an equilibrium state is reached and the fraction of looped DNA remains almost constant. The fraction of DNA constructs that had formed loops ($f[L]$) is determined and plotted as a function of Smc5/6 concentration ($[L]$) and fitted with the Hill-Langmuir function: $f([L]) = [L]^n/((K_a)^n + [L]^n)$, where $K_a$ is the concentration of Smc5/6 at which half of the DNA is looped and $n$ is the Hill-coefficient. DNA molecules whose end-to-end distance was larger than 10 µm were not counted for this analysis as they are unlikely to act as DNA substrates for loop extrusion due to the high tension/stall force on the stretched DNA (Fig. 1M, Fig. S5B).

**Probability of bleaching steps from dimer to monomer ratio.**

We determine the probability P(N) of observing either N=0, 1, or 2 bleaching steps as function of the dimer fraction x we use the following formulas:

$$P(2) = p^2x,\ P(1) = 2(1-p)px + px,\ P(0) = (1-p)(1-x) + (1-p)^2x,$$

where p indicates the labeling efficiency. The respective errors are calculated using: $\sigma(P(N)) = \pm\frac{dP(N)}{dp}\sigma_p$ , where $\sigma_p$ is the error of the labeling efficiency.

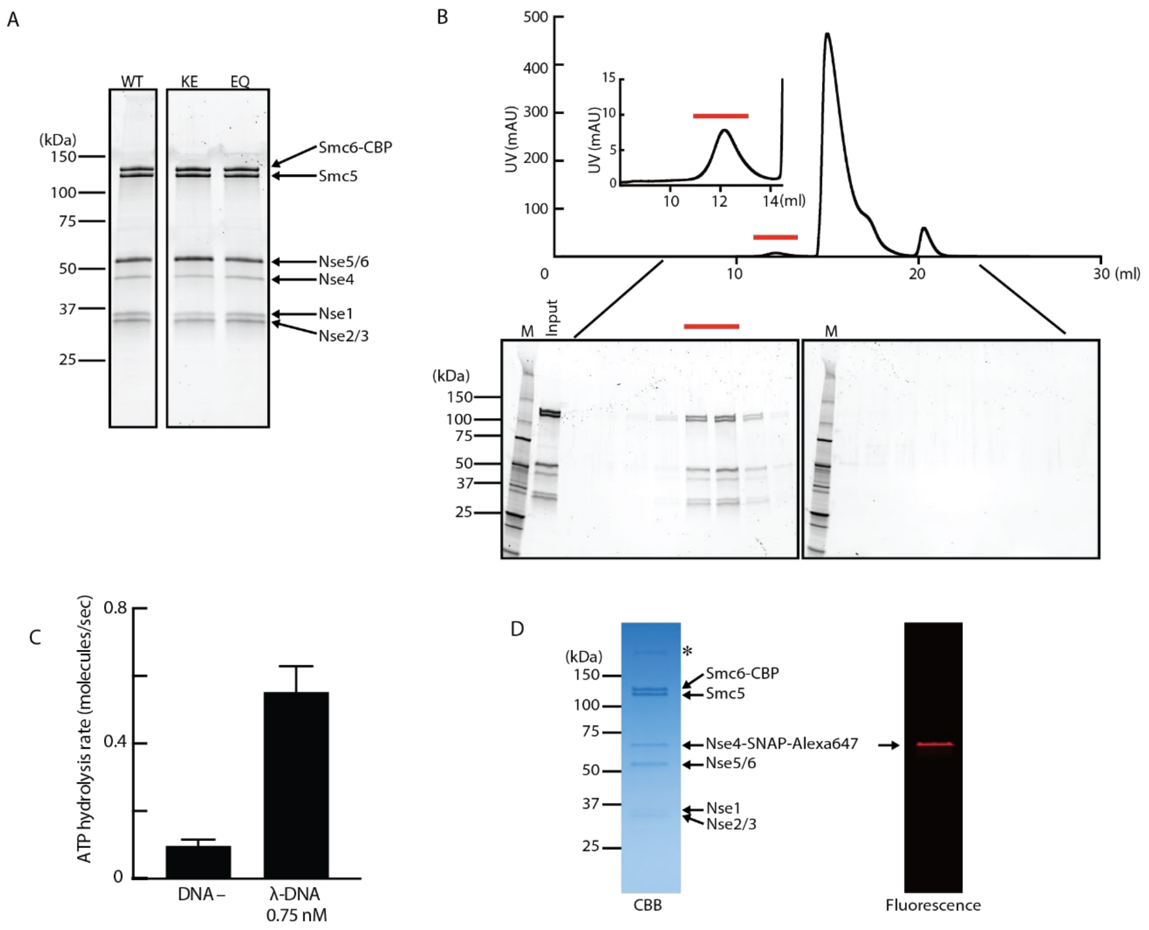


**Fig. S1. Purification and ATPase activity of the Smc5/6 complex.** (A) Oriole stained SDS-PAGE gel for purified Smc5/6 and mutants. (B) The result of size-exclusion chromatography (SEC) of wild type Smc5/6. The chromatogram and Oriole stained SDS-PAGE for each fraction are shown. Peak fractions are indicated with red bar. (C) ATP hydrolysis rate for wild type Smc5/6 with/without λ-DNA. (D) Fluorescently labeled Smc5/6. CBB staining and fluorescence detection of SDS-PAGE are shown.

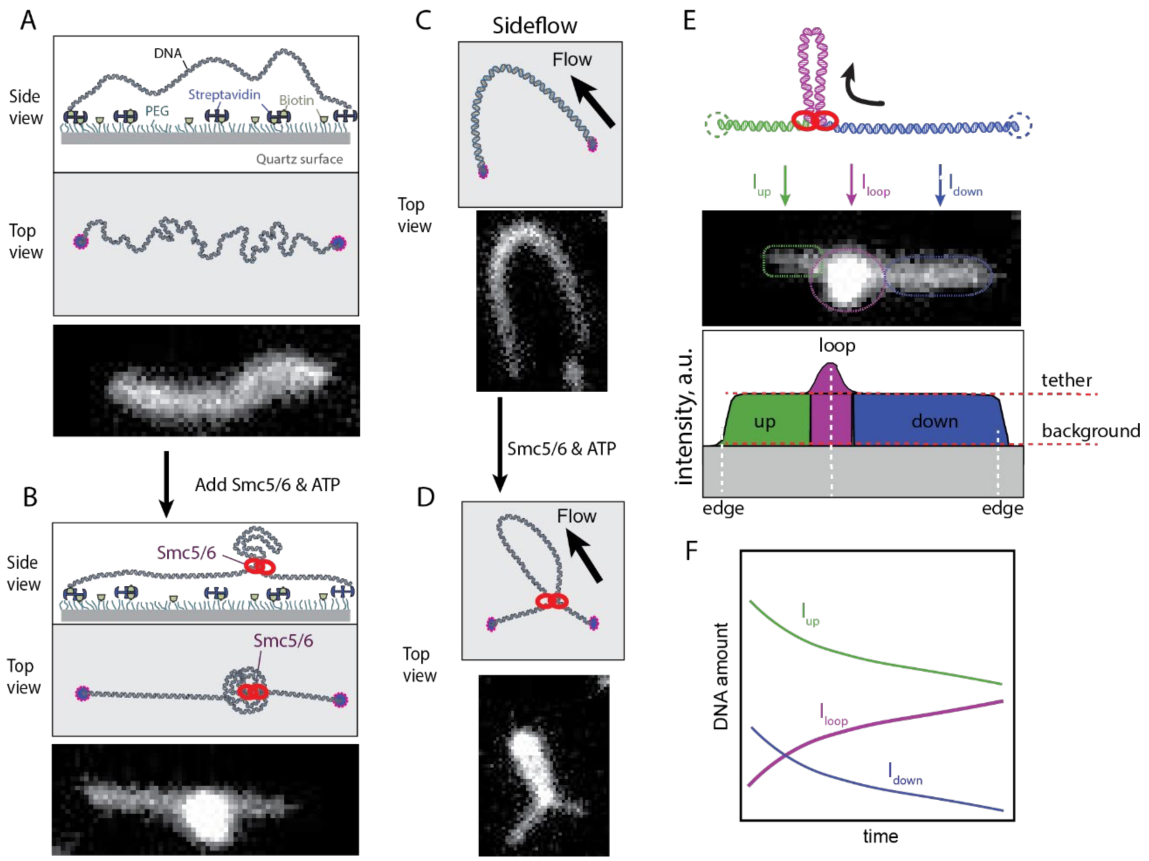


**Fig. S2. Schematic of the loop extrusion assay and analysis.** (A) Side-(top) and top-view (center) of phage λ-DNA anchored onto glass substrate passivated with PEG molecules alongside an experimental snapshot of the DNA from the top-view (bottom). (B) Schematics showing side- (top) and top-view (center) of DNA condensing into a punctum upon addition of Smc5/6 and ATP. The bottom plot shows an experimental snapshot. (C) Schematic (top) and experimental snapshot (bottom) showing an arc-shaped DNA due to buffer application. (D) Schematic (top) and experimental snapshot (bottom) showing a typical DNA loop extruded from a specific point along the DNA strand under sideway flow. (E) Schematic (top), snapshot (middle), and a schematic of fluorescence intensity profile of the DNA molecule with a loop, showing the 3 different sections ($I_{up}$, $I_{loop}$, $I_{down}$) (F) Schematic showing the DNA length changes calculated from the integrated fluorescence intensities for the three different zones ($I_{up}$, $I_{loop}$, $I_{down}$) plotted with time.

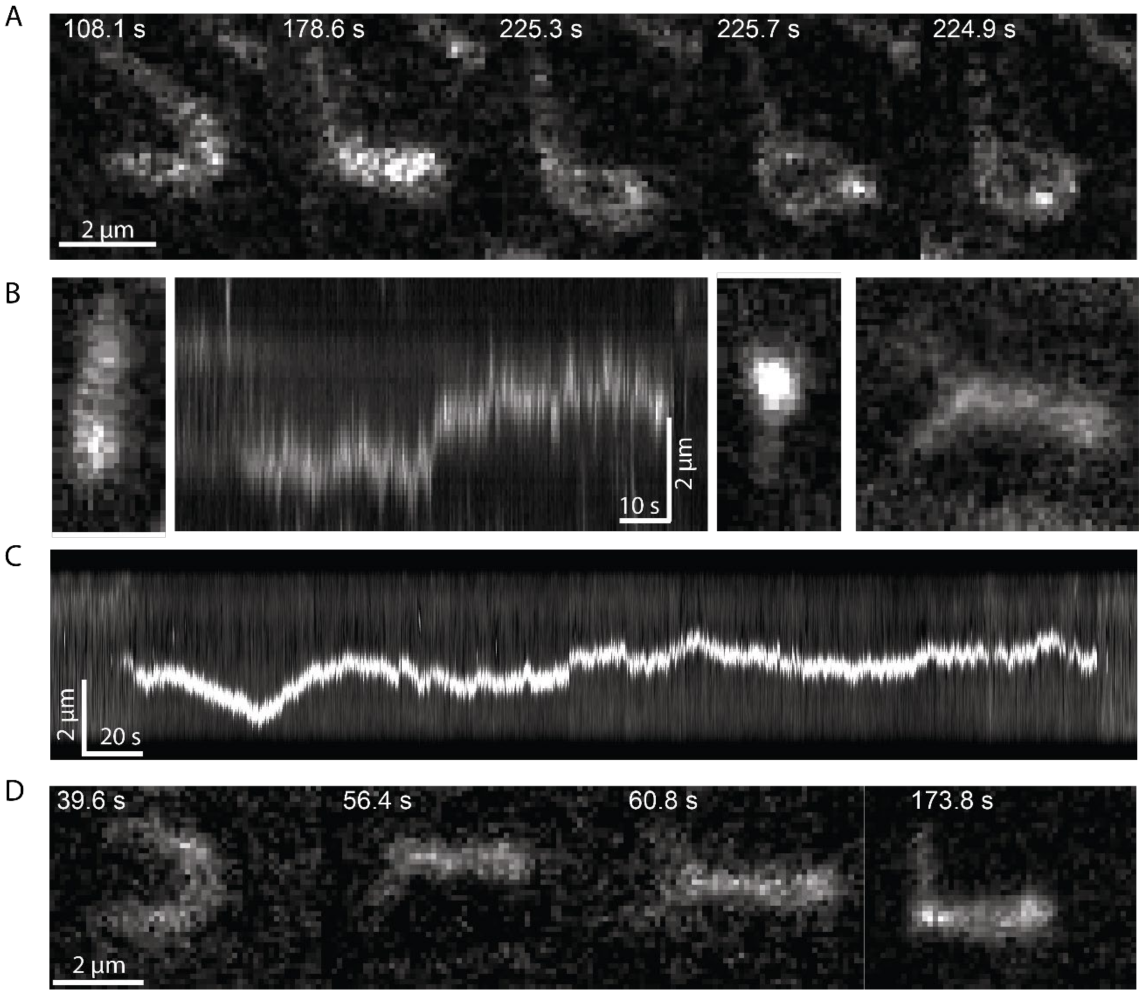


**Fig. S3. Characteristics of the Smc5/6-mediated DNA loop extrusion.** (A) Snapshots showing the process of loop extrusion on a sticky surface under side flow revealing the O-shape topology of the DNA loop. (B) First to third plot: Snapshots (first and third, from the left) from the first and the last frames of the kymograph (second) showing a loop extrusion event in the absence of flow. The fourth plot shows the same loop upon the application of side-flow confirming that the observed DNA punctum is a loop. (C) A kymograph of a DNA molecule showing a Smc5/6-mediated loop that exhibits diffusion along the DNA. (D) Snapshots showing diffusion of an extruded loop. The side-flow reveals that the stem of the loop moves along the DNA.

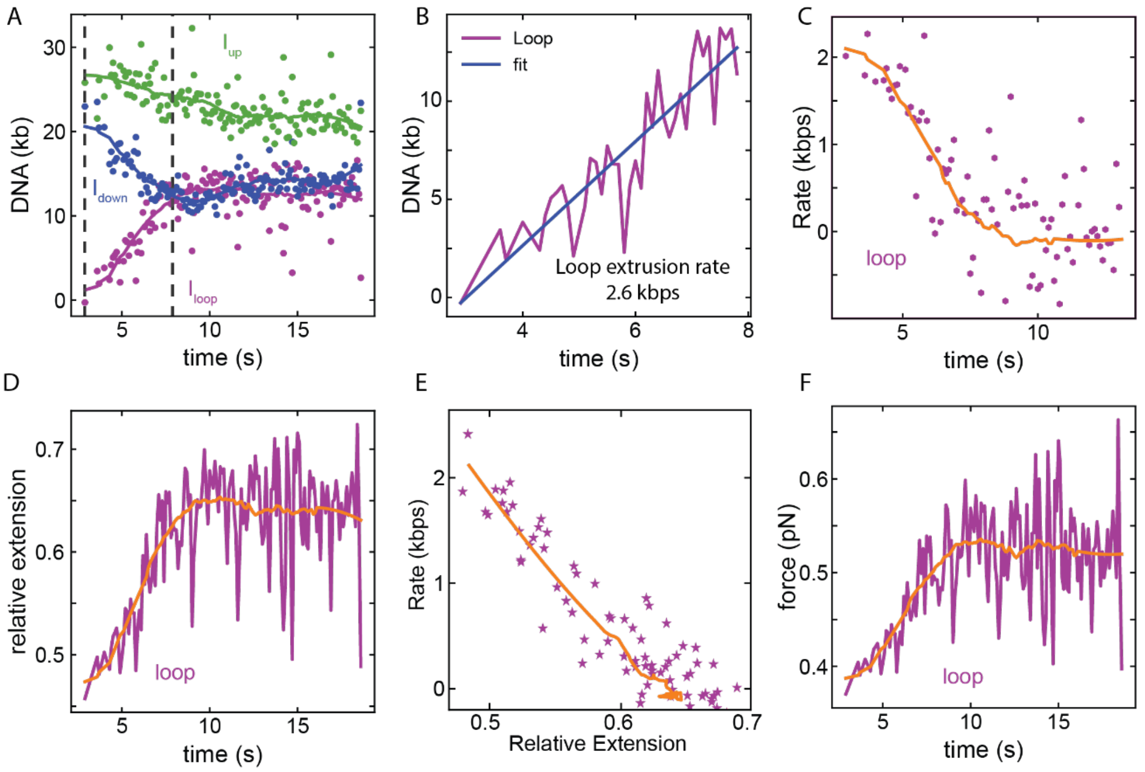


**Fig. S4. Quantitation of Smc5/6-mediated loop extrusion kinetics extracted from a single looping event.** (A) Kinetics of loop extrusion showing changes in DNA sizes from different sections ($I_{down}$, $I_{loop}$, $I_{up}$) over time. The linear region within the two dashed lines was used to obtain the initial loop extrusion rate via linear fit as shown in. (B). (C) The change of loop extrusion rate during the loop growth, which was calculated using the change in loop sizes in a moving time window of 2 s. (D) The simultaneous change in relative DNA extension as a function of time. (C, D) shows that the decrease of extrusion rate is correlated with the increase in DNA tension. (E) Scatter plot of loop extrusion rate with the relative extension, taken from (C, D), showing that rate drops nearly zero above extension value of 0.6. (F) Change in tension on the DNA with time. The force values were extracted from the values of relative DNA extension in (D) and converted to the known force–extension relation. (A-F) corresponds to the single DNA loop extrusion shown in Fig. 1F-H, and solid lines show running averages over 50 points.

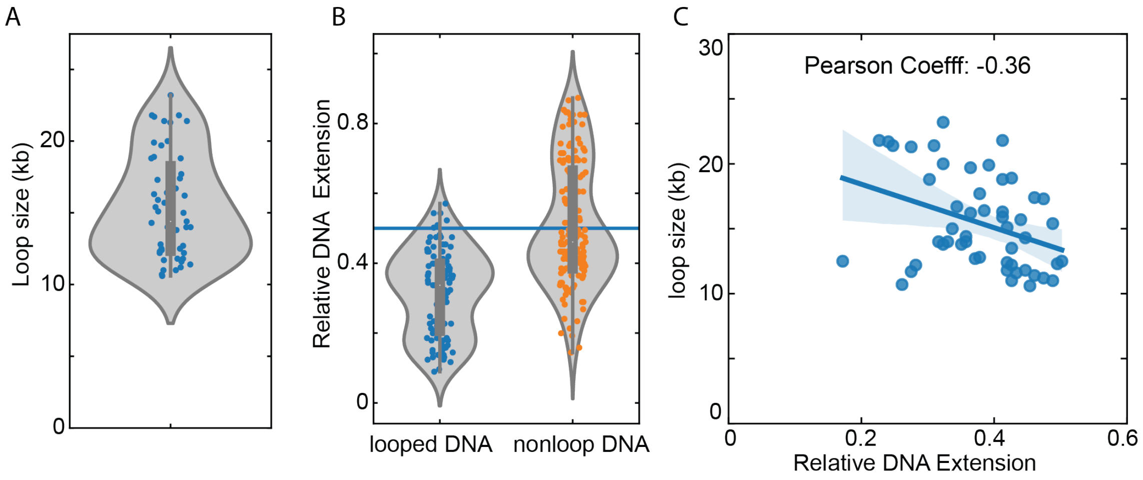


**Fig. S5. Statistics of the characteristics of Smc5/6 extruded DNA loops.** (A) Loop size distribution (violin plot) from 100 different DNA looping events. (B) Violin plot distributions of relative extension of DNA which formed loops (N=100) and did not form loops (orange, N=140). Extension values were taken before the start of loop extrusion event. Only 6% of the loop extrusion events (N=100) were observed on DNA molecules which were stretched beyond an extension value of 0.6 (indicated by the horizontal line). (C) Scatter plot of loop sizes versus the relative DNA extension showing a negative correlation with Pearson coefficient of -0.36. The line shows a linear regression fit with 95 % confidence interval indicated as shaded area.

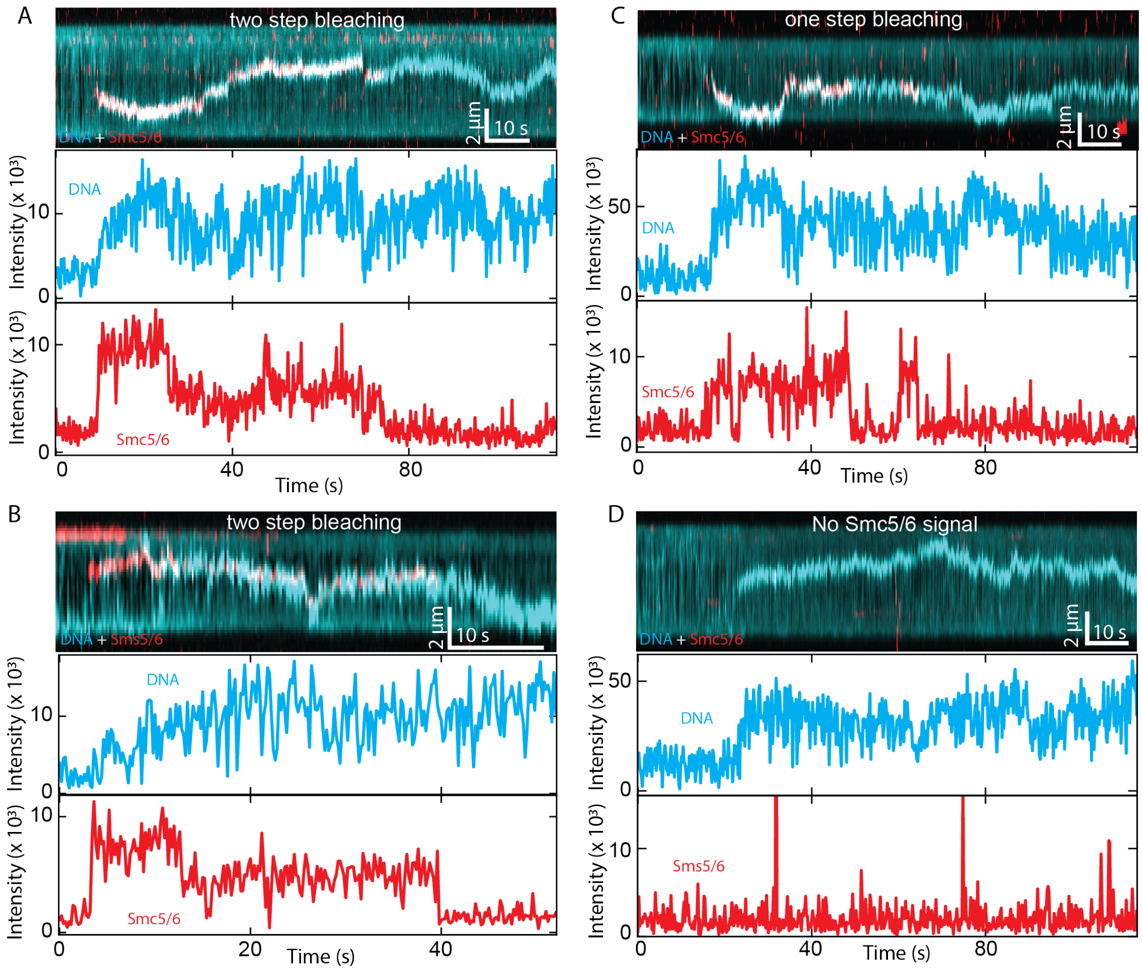


**Fig. S6. Additional examples of photobleaching events from Alexa-647 labeled Smc5/6 during DNA loop extrusion.** Kymograph (top of each section) of DNA (cyan) and Smc5/6 (red). Time traces of DNA (middle of each section) and Smc5/6 fluorescence intensity (bottom of each section) determined from the corresponding kymographs showing two bleaching steps (A, B), one bleaching step (C), and no Smc5/6 signal at the loop initiation position (D). Intensity values displayed in Smc5/6 time traces were extracted on the same pixel positions as the DNA puncta intensities.

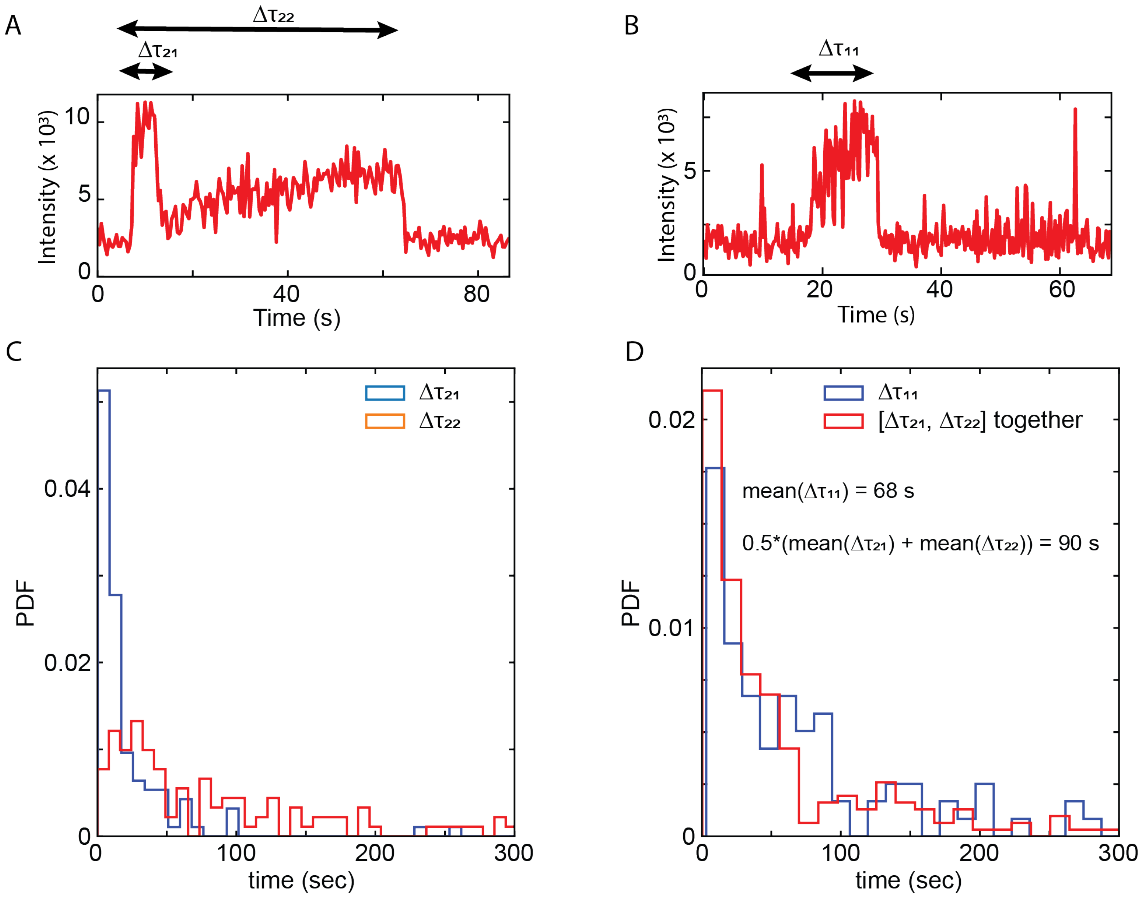


**Fig. S7 Bleaching time distribution.** (A, B): Schematics showing how the bleaching times $\Delta\tau_{21}$ and $\Delta\tau_{22}$ (A), and $\Delta\tau_{11}$ (B) are determined. (B) The bleaching time $\Delta\tau_{11}$ in a one-step bleaching trace and corresponding distribution shown in D (blue). When the two bleaching times $\Delta\tau_{21}$, $\Delta\tau_{22}$ from the two-step bleaching trace are put together in a histogram, they resemble the single-step distribution. Note that the average time of one-step bleaching traces is similar to the average of the times ($\Delta\tau_{21}$, $\Delta\tau_{22}$) of the two step bleaching traces.

A

B

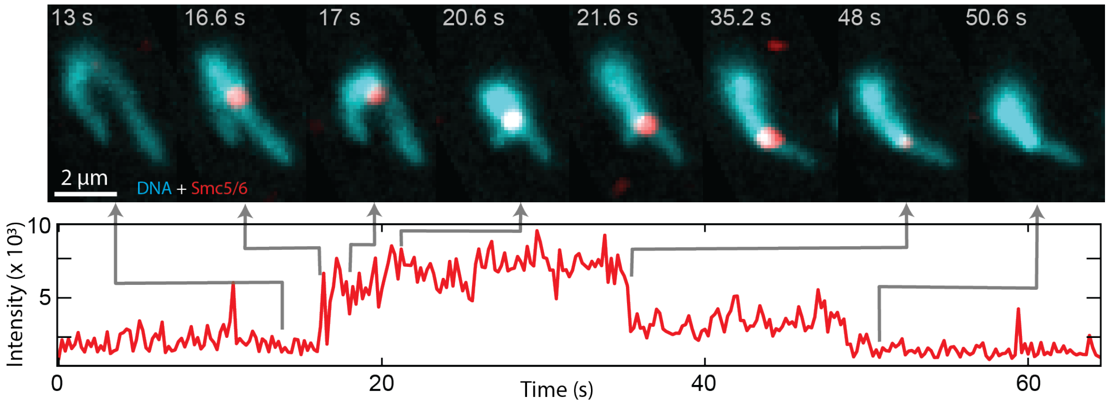


**Fig. S8. Two bleaching steps observed from labeled Smc5/6 at the stem of the DNA loop during loop extrusion.** (A) Snapshots of DNA loop extrusion event showing labeled Smc5/6 at the stem of the extruded loop. (B) the corresponding time trace Smc5/6 fluorescence intensity showing two-step bleaching. The arrows indicate the time points corresponding to the respective snapshots.

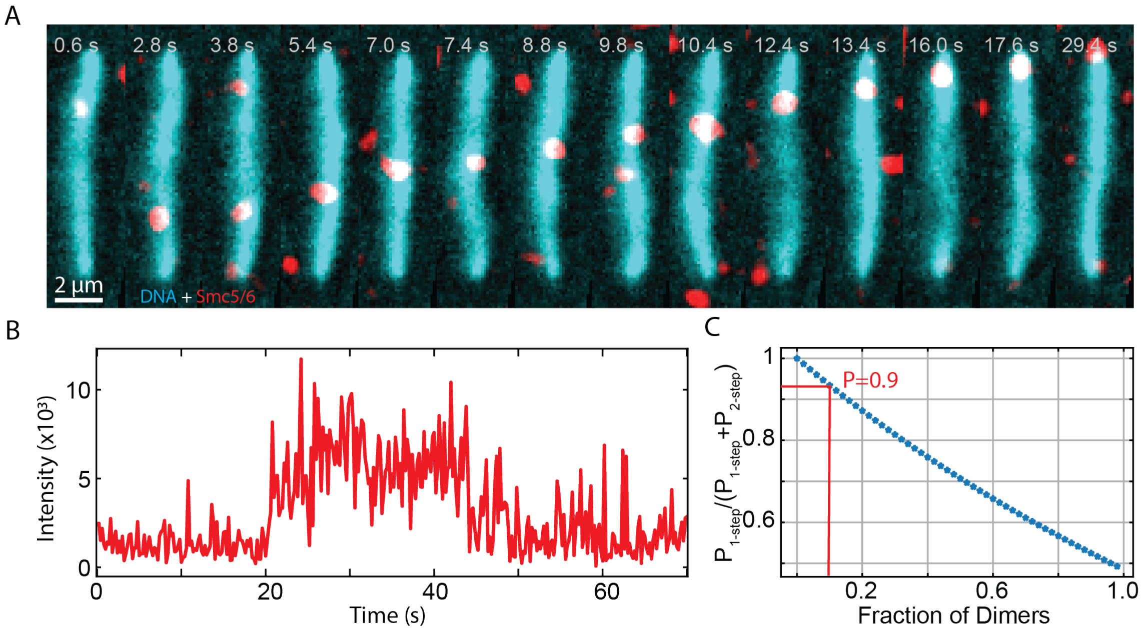


**Fig. S9. DNA Translocation by a single Smc5/6 complex.** (A) Snapshots of labelled Smc5/6 (red) on DNA (cyan), showing a DNA translocation event, corresponding to Fig.3A. (B) Intensity time trace of the Smc5/6 depicted in (A) showing one-step bleaching. (C) The calculated probability of observing single bleaching steps as a function of dimer fraction given that we experimentally only observe one-step and two-step bleaching events.

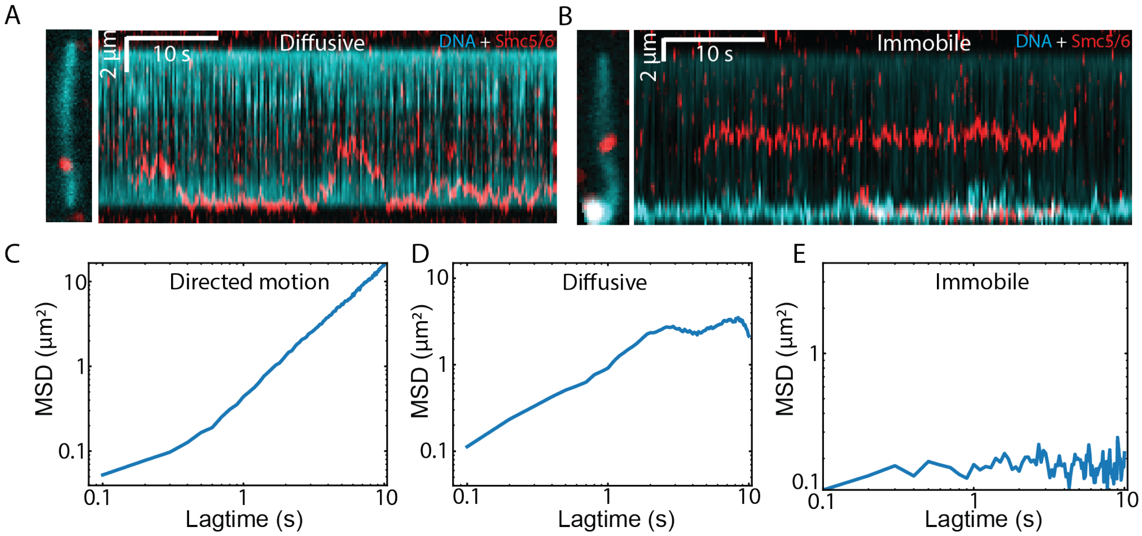


**Fig. S10. Different not loop-extruding interaction modes between DNA and Smc5/6.** Example kymographs of Smc5/6 (red) on DNA (cyan) showing diffusive (A) and immobile characteristics (B). (C, D, E) Mean square displacements (MSD) of Smc5/6 complexes displaying directed motion (C) corresponding to Fig. 3A, diffusive motion (D) corresponding to Fig. S10A, and immobile behaviour (E) corresponding to Fig. S10B.

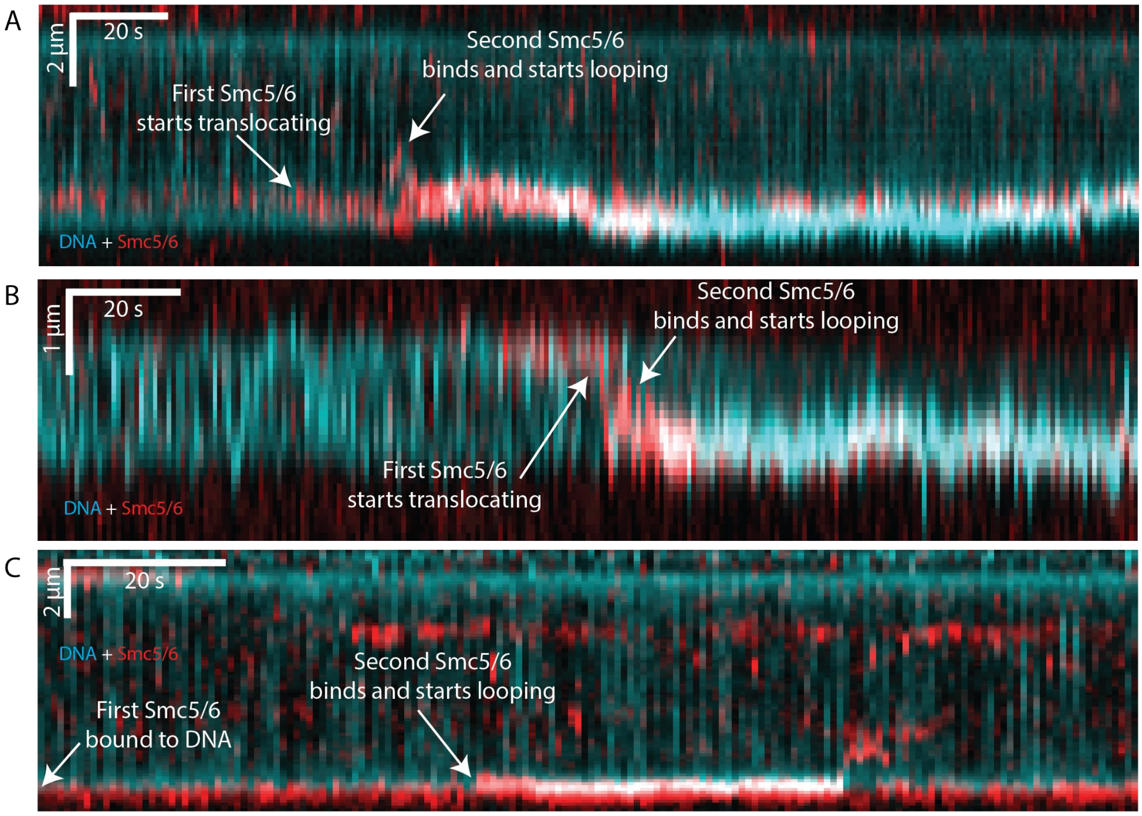


**Fig. S11. Examples showing the cases where dimerization of Smc5/6 result in initiation of loop extrusion.** Example kymographs showing different types of events where loop extrusion (cyan) starts upon dimerization of Smc5/6 (red). (A) Two translocating Smc5/6 merge together and start loop extrusion. (B) A single Smc5/6 bound on the end of DNA starts translocating. The second Smc5/6 from the solution binds the first one to initiate loop extrusion. (C) A single Smc5/6 is bound on one end of the DNA at the start of the measurement. The second Smc5/6 binds the first one from the solution to initiate loop extrusion.

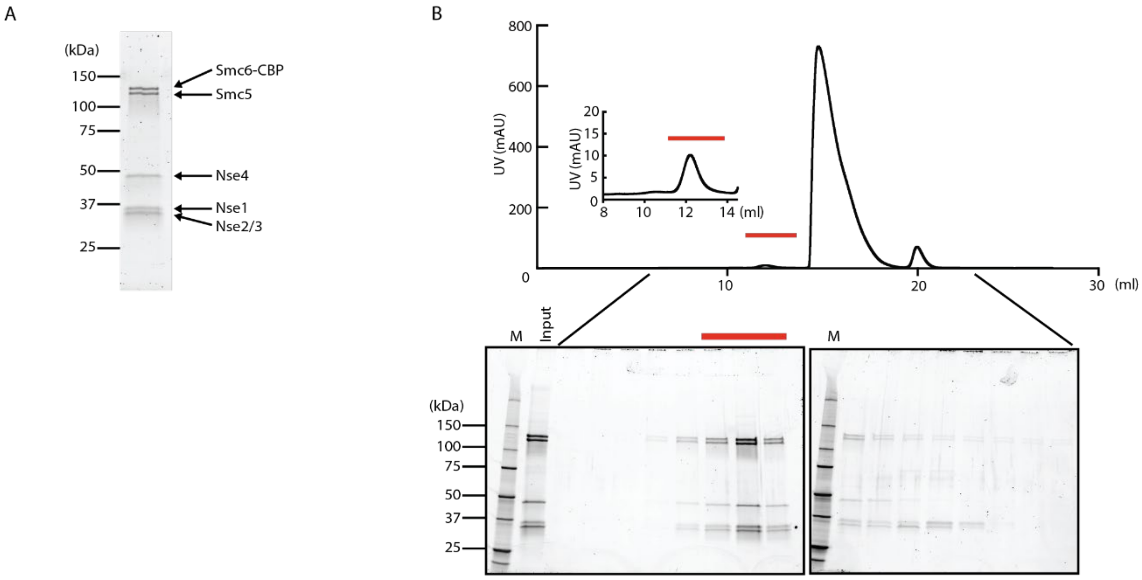


**Fig. S12. Purification of hexameric Smc5/6 complex.** (A) Oriole stained SDS-PAGE for the Smc5/6 complex lacking Nse5/6 (hexamer) (B) The result of SEC for the hexamer. The chromatogram and Oriole stained SDS-PAGE for each fraction are shown. Peak fractions are indicated with red bar.

**Captions for Supplementary Videos**

**Movie S1.** Side-flow visualization of DNA loop extrusion by Smc5/6 corresponding to Fig. 1E.

**Movie S2.** DNA loop extrusion by Smc5/6 without side-flow along with the kymograph of the DNA.

**Movie S3.** Side-flow visualization of DNA (cyan) loop extrusion by labelled Smc5/6 (red) where Smc5/6 appears on the DNA at ~7s followed by loop extrusion. The Smc5/6 stays at the stem of the loop before finally bleaching at ~151 s.

**Movie S4.** DNA loop extrusion by labelled Smc5/6 without side-flow. The top kymograph and image sequences correspond to DNA and the middle section corresponds to the labelled Smc5/6. The merged kymograph and image sequences of DNA and Smc5/6 can be seen in the bottom lane.

**Movie S5.** Directional translocation of a Smc5/6 (red) on the DNA (cyan). The left panel is the kymograph corresponding to the time lapse video of DNA and Smc5/6 on the right panel.

**Movie S6.** A single translocating Smc5/6 complex dimerizes with another Smc5/6 (red) to initiate DNA (cyan) loop extrusion.
